# Change in brain molecular landscapes following electrical stimulation of the nucleus accumbens

**DOI:** 10.1101/2024.09.30.615737

**Authors:** Chengwei Cai, Lingyun Gao, Zhoule Zhu, Wangyu Chen, Fang Zhang, Chaonan Yu, Kedi Xu, Junming Zhu, Hemmings Wu

## Abstract

Deep brain stimulation (DBS) targeting the nucleus accumbens (NAc) is a promising therapeutic intervention for treatment-resistant neuropsychiatric disorders such as depression, anxiety, and addiction. However, the molecular mechanisms underlying the clinical efficacy of NAc DBS remain largely unknown. One approach to address this question is by performing spatial gene expression analysis on cells located in different regions of the same circuit following NAc DBS. In this study, we utilized high-resolution spatial transcriptomics (Stereo-seq) to investigate gene expression changes induced by NAc DBS in the mouse brain. Mice were randomly allocated to received continuous electrical stimulation (0.1 mA, 130 Hz) or sham treatment (electrode implanted, no electrical stimulation given) for one week, and subsequent Stereo-seq analysis identified differentially expressed genes (DEGs) across various brain regions. Our findings reveal widespread alterations in synaptic function and neuronal connectivity, particularly in glutamate receptor-expressing neurons in the NAc, which play a key role in the reward circuitry. Functional enrichment analysis highlighted changes in transporter activity and glutamate receptor binding in brain regions such as the anterior cingulate cortex and lateral septal nucleus. Single-cell resolution mapping further identified key molecular players, including Nlgn1, Snca, PDE10a, and Syt1, which are critical for synaptic plasticity and neurotransmitter release, and have been implicated in various psychiatric disorders. These findings shed light on the molecular underpinnings of NAc DBS and provide insights into its therapeutic potential in modulating neural circuits associated with neuropsychiatric disorders.

## INTRODUCTION

Reward circuitry dysfunction is highly implicated in multiple neuropsychiatric disorders, including major depressive disorder, anxiety disorder, obsessive-compulsive disorder, eating disorder, and addiction. At the center of the reward circuitry, which plays a crucial role in the processing of pleasure and reinforcement learning, lies the nucleus accumbens (NAc), a key structure involved in the experience of reward and motivation. The main inputs to NAc include dopaminergic projections from the ventral tegmental area, and glutamatergic projections from the lateral hypothalamus, hippocampus and prefrontal cortex^1-4^. Selective manipulations of the NAc neural pathways have shown to reduce depression-, anxiety-, compulsion-, anorexia-, and addiction-like phenotypes in animal models^5-8^. Neuromodulation targeting the NAc has also been proposed as a potential clinical treatment for these neuropsychiatric disorders^9^.

Deep brain stimulation (DBS) is a neuromodulation technique, in which high-frequency (usually between 100 to 200 Hz) electrical stimulation is delivered through electrodes implanted in specific brain regions. Clinical trials using DBS targeting the NAc region have shown preliminary but promising results for a number of otherwise-refractory neuropsychiatric disorders, including depression, anxiety, obsessive-compulsive disorder, anorexia nervosa, and addiction^10-13^.

The most widely accepted theory concerning the action mechanism of DBS suggests that the procedure disrupts pathological neural pathways and restores normal functions through the delivery of artificially generated electrical impulses^14, 15^. This neuromodulatory intervention is thought to modulate neural circuits by changing the activity of neurons within the targeted region, as well as those in connected regions, thereby rectifying the dysfunctional signaling associated with neuropsychiatric conditions. Extensive research into the action mechanism of DBS reveals that high-frequency stimulation may lead to both immediate and long-term neuromodulatory effects. Immediate effects encompass the depolarization blockade of neuronal activity, synaptic inhibition, and synaptic potentiation, which can alter the firing patterns of neurons in the NAc and its afferent and efferent connections^16-18^. Over the long term, DBS has been shown to induce neuroplastic changes, associated with the reorganization of synaptic connections, potentially underpinning the sustained therapeutic benefits observed in patients^19-21^. Moreover, investigations have indicated that DBS may exert its effects by modulating neurotransmitter release, such as dopamine, glutamate, and GABA, which are integral to mood, motivation, and reward processes. For instance, increased dopaminergic transmission following NAc DBS may contribute to enhanced mood and motivation in patients with depression^22^. Similarly, the normalization of glutamate and GABA levels may help alleviate symptoms of anxiety and obsessive-compulsive disorder^23, 24^. In addition to neurotransmitter systems, DBS has been linked to changes in gene expression and the upregulation of neurotrophic factors, supporting neuronal survival, growth, and synaptic plasticity^25, 26^. These molecular alterations could be part of the mechanism by which DBS induces long-lasting modifications in neural networks and cognitive and emotional processes.

Despite these insights, the exact cellular and molecular pathways through which NAc DBS exerts its therapeutic effects remain only partially understood. The various responses observed across different disorders suggest that DBS may influence multiple molecular targets within the reward circuitry. However, the current understanding for the action mechanisms of NAc DBS at cellular and molecular levels remain confined, partly due to technological limitations.

Spatial transcriptomics, a novel technology first described in 2016, has empowered researchers to visualize and map the gene expression patterns of individual cells within their spatial context in a tissue sample, thereby providing a comprehensive view of the cellular and molecular complexity of tissues in a spatially resolved manner^27^. It has been used to study the molecular changes during normal brain development, maturation, and aging, as well as in disease models^28^. Here, we describe the use of spatial transcriptomics to investigate the differentially expressed genes at accumbal, network, and global levels following NAc DBS, in hope of revealing the molecular mechanisms of action. Our inquiries are twofold: First, to determine the general effect of NAc DBS on global gene expression, and second, to elucidate the specific network effect of NAc DBS on gene expression.

To address these questions, our study employed a multidisciplinary approach, combining neuroanatomical techniques with spatial transcriptomics to map the NAc’s transcriptomic landscape post-DBS. The results of this study are expected to provide a molecular blueprint for the therapeutic effects of NAc DBS, and pave the way for refined neuromodulatory strategies targeting the NAc in the treatment of neuropsychiatric disorders.

## METHODS AND MATERIALS

### Animals

C57BL/6J male mice (6–8 weeks of age, n = 31) were obtained from the Shanghai SLAC Laboratory Animal Co., Ltd. Mice were socially housed in the animal facility under standard condition (12h light/dark cycles), and were allowed to acclimatize to their environment for 2 weeks before experimental procedures. All animal experiments were approved by the Animal Care and Use Committee of the animal core facility at Zhejiang University School of Medicine (Approval No. AIRB-2021-087).

### Surgical Procedures

After accommodation, mice were implanted with bipolar platinum electrode (Kedou (Suzhou) Brain Computer Technology, KD-BSEC) in the left NAc (0.8 mm anterior to Bregma, 1.3 mm lateral to midline, and 4.25 mm deep at 90 degree angle from the brain surface). After surgery, the animals were single-housed to recover for 1 week. Then the mice were assigned randomly into experimental group (DBS, n = 20) or control group (n = 11), which would receive continuous electrical stimulation for 7 days (0.1 mA, 130 Hz, 100 μs, bipolar, biphasic) and sham stimulation for 7 days, respectively.

### Sample preparation

After stimulation, mice were euthanized with sodium pentobarbital (450 mg/kg intraperitoneally). Brain samples were collected and embedded immediately with Tissue-Teck OCT. Coronal sections of samples were macroscopically examined in a Leica CM1950 cryostat (Buffalo Grove, IL, USA) until electrode tracts were reached. Hemispheres with electrodes best positioned in the NAc (DBS: n = 2, control: n =2) were selected, and coronal sections (10 *µ*m thick) were performed to obtain the final section for Stereo-seq.

### Stereo-seq library construction and sequencing

The Stereo-seq library preparation and sequencing was performed as previously described^29^. In brief, brain sections were fixed in pre-chilled methanol (−20°C, 40 min), then permeabilized on chips with 100 mL of 0.1% pepsin in 0.01 M HCl (37°C, 5 min). Permeabilization solution was then removed and sections were rinsed with 100 mL 0.1 SSC buffer (Thermo, CA, USA). The mRNAs captured by DNA nanoballs (DNBs) on the chip were then reverse transcribed by SuperScript II reverse transcription (RT) mix (80 *µ*l RT Reagent, 5 *µ*l Reverse T Enzyme, 5 *µ*l RI, 5 *µ*l RT Oligo, 5 *µ*L RT Additive, a total of 100 *µ*l mix per chip, BGI, Shenzhen, China) at 42°C for 1.5 h. After removing the RT mix, the chip was washed with 0.1X SSC (Thermo, CA, USA) and incubated at 37°C for 30 min in TR Buffer (BGI, Shenzhen, China). Then the chips were washed twice with 0.1X SSC (Thermo, CA, USA), and immersed in the cDNA Release Mix (380 *µ*l cDNA Release Buffer, 20 *µ*l cDNA Release Enzyme, a total of 400 *µ*l mix per chip, BGI, Shenzhen, China) for 3 h at 55°C. The resulting cDNAs were amplified with PCR Mix (42 *µ*l Eluted cDNA, 8 *µ*l cDNA Primer, 50 *µ*l cDNA Amplification Mix, a total of 100 *µ*l mix per reaction, BGI, Shenzhen, China). After PCR, the mix was washed 10 times with nuclease-free H_2_O (Thermo, CA, USA) and was purified using the Ampure XP Beads (Vazyme, Nanjing, China). The resulting cDNAs eluted by nuclease-free H_2_O were amplified with PCR Mix (50 *µ*l cDNA Amplification Mix, 42 *µ*l Eluted cDNA, 8 *µ*l cDNA Primer, a total of 100 *µ*l mix per reaction, BGI, Shenzhen, China). The purified cDNA products were then fragmented by Fragmentation Mix (4 *µ*l TMB, 1 *µ*l 10-fold diluted TME, 15-x *µ*l nuclease-free water, x *µ*l cDNA product, a total of 20 *µ*l mix per reaction, BGI, Shenzhen, China) at 55°C for 10 min. After fragmentation, the products were amplified with PCR Mix (50 *µ*l PCR Amplification, 25 *µ*l PCR Barcode Primer Mix, 25 *µ*l Fragmentation Mix, a total of 100 *µ*l mix per reaction, BGI, Shenzhen, China). The purified cDNA mix was used for DNB generation. Sequencing of the final libraries was performed on a MGI DNBSEQ-Tx.

### Raw data processing

In this study, raw sequencing data was processed using the SAW pipeline^30^. In brief, we first extracted the unique molecular identifiers (UMI, bases 26-35 of read1), and cDNA (whole reads 2) and coordinate identity (CID, first 25 bases of read1) from the paired-end sequencing data. These were then mapped to a pre-position-resolved CID whitelist using the ST_BarcodeMap software (https://github.com/BGIResearch/ST_BarcodeMap), which allowed for a tolerance of 1 mismatched base. Subsequently, any read pairs lacking valid CID sequencing data were discarded. We further refined the dataset by removing read pairs with low-quality UMIs, identified either by the presence of any undefined (N) bases or by having more than 2 bases with a quality score under 10 (Phred + 33). The cDNA sequences with valid CID and UMI were then aligned to the mm10 mouse reference genome using STAR software^31^. Finally, we excluded alignments that exhibited a mapping quality score (MAPQ) less than 10, and annotated the filtered alignments to the corresponding genes.

### Binning data acquisition

DNB images were first manually registered with nucleic acid staining images. The single-cell segmentation was then performed using Scikitimage after removing the background and computing Euclidean distances^32^. Transcripts captured by 100*100 DNBs were merged as one bin 100, which we treated as the fundamental analysis unit. The bin IDs were derived from their spatial coordination (spatial_x and spatial_y) at the capture chip.

### Single-cell data acquisition

Based on the bin 1 size matrix file, the single-cell gene expression matrix was generated for downstream analysis by summarizing all UMIs from DNBs within a putative single cell. The X-Y coordinates of each cell and the overall MID count were obtained using Stereopy^33^ and processed for subsequent analysis.

### Segmentation of the region of interest

Brain regions of interest, including the cingulate cortex (CC), caudate putamen (CPu), meninges, cortex, intermediate part of the lateral septal nucleus (LSI), nucleus accumbens core (NAc.c), nucleus accumbens shell (NAc.s), anterior part of the anterior commissure, and nucleus of the vertical limb of the diagonal band (VDB) were automatically segmented using Stereopy^33^. Other brain regions of interest, including anterior cingulate cortex (ACC) and claustrum (CL), were manually segmented based on spatial map of UMI count of each sample, and the corresponding bin 1 matrix was extracted.

### Quality control

Processing of the MID count matrix obtained from Stereo-seq bin 100 data was implemented using the R package Seurat (v4.2.2)^34^. To remove low-quality cells, we filtered out cells with gene numbers no more than 200 or higher than 5000. In addition, we removed cells with high mitochondrial genes (>10%) to avoid perforated cells that lost cytoplasmic RNA. Mitochondrial reads were subsequently removed, and Genes *Gm42418* and *AY036118* were also removed, as they overlapped the rRNA element Rn45s and represented rRNA contamination.

### Unsupervised clustering and annotation

Seurat was used for quality control, normalization, dimensionality reduction, clustering, and identification of marker genes^34^. Sections from different samples were integrated by the R package harmony^35^. Brain region definition was achieved by combining morphological information and spatially constrained clustering. The identification of cell types was based on the known marker gene sets, and verified by in situ hybridization results from the Allen Brain Atlas (https://atlas.brain-map.org/).

### DEG analysis

Genes differentially expressed among cell types (or cell subtypes) were identified by comparing average transcript expressing levels in cells of a given cell type (or cell subtypes) between groups using the Wilcoxon Rank Sum test in Seurat package. For cell types, the default parameters were used. For cell subtypes, the following cutoffs were applied: logfc.threshold = 0.2, min.pct = 0.1.

### Gene Ontology enrichment analysis

R package clusterProfiler was used to identify enriched Gene Ontology terms^36^. Marker gene lists from indicated cell types were inputted with default parameters.

## RESULTS

### Stereo-seq profiles spatial transcriptomes of mouse brain

To fully characterize the spatial transcriptomes alterations induced by NAc DBS, we utilized the Stereo-seq (spatial enhanced resolution omics sequencing; Fig. 1A). Expression profile matrix was first binned into bin 100, and subsequently processed by quality control, batch correction and unsupervised clustering, after which a total of 15 clusters with 27,241 bins were obtained (Supplementary Fig. S1A-D). The clusters were mapped to the spatial atlas according to the coordinate information acquired by the CID label (Fig. 1A,B). Based on anatomical structures and canonical markers, we annotated the clusters with 9 structures, including the cingulate cortex (CC), caudate putamen (CPu), meninges, cortex, intermediate part of the lateral septal nucleus (LSI), nucleus accumbens core (NAc.c), nucleus accumbens shell (NAc.s), anterior part of the anterior commissure, and nucleus of the vertical limb of the diagonal band (VDB). To ensure the reliability of the data, we further compared the spatial distribution of Stereo-seq signals of selected marker genes with in situ hybridization results from the Allen Brain Atlas (Fig. 1C). Overall, the Stereo-seq data revealed structural composition of brain tissues, facilitating reliable downstream spatial transcriptomic analysis with specific regional gene expression profiles.

**Fig. 1.**
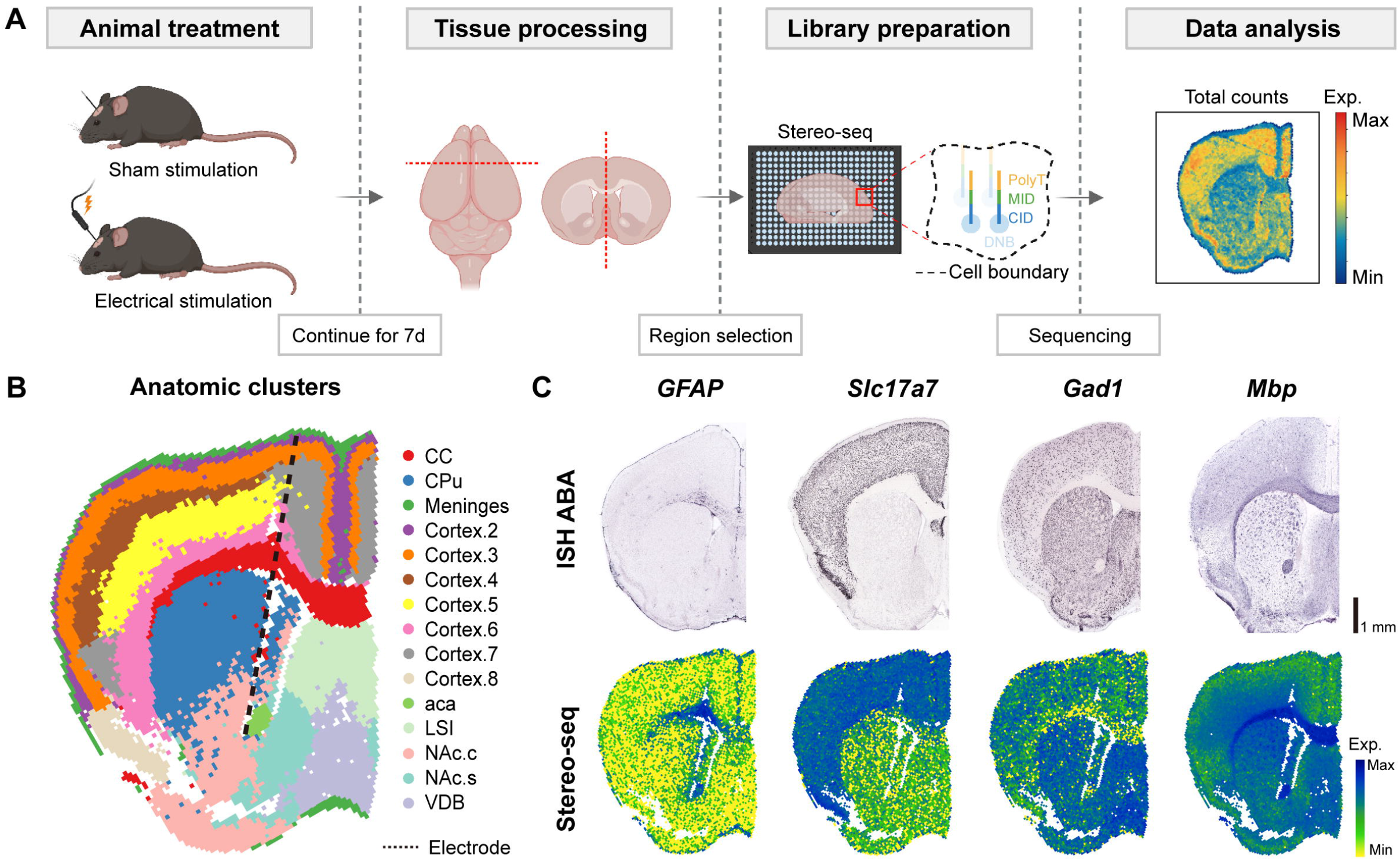
Stereo-seq captures spatially resolved hemispheric gene expression **after electrode implantation in the nucleus accumbens**. (**A**) Overview of the experimental workflow. Continuous electrical or sham stimulation was applied to the nucleus accumbens (NAc) for 7 days, followed by tissue processing, library preparation, and spatial transcriptomic data analysis. (**B**) Spatial transcriptomic analysis using Stereo-seq. Several anatomic clusters were identified, including cortical regions and subcortical areas. Specific regions include the cingulate cortex (CC), caudate putamen (CPu), meninges, cortex, lateral septal nucleus (LSI), nucleus accumbens core (NAc.c), nucleus accumbens shell (NAc.s), anterior part of the anterior commissure (aca), and nucleus of the vertical limb of the diagonal band (VDB). (**C**) Gene expression verification across hemispheres. The spatial distribution of genes such as GFAP, Slc17a7, Gad1, and Mbp was mapped across various brain regions. Comparisons were made between sham-stimulated samples and data from the In Situ Hybridization Allen Brain Atlas (ISH ABA), showing aligned gene expression distributions. Scale bar, 1 mm. CID: coordinate identifier, DNB: DNA nanoball, MID: molecular identifier. Panel A was created using BioRender platform (www.biorender.com).

### NAc DBS drives global alteration of gene expression

In order to understand the electrogenetic modulatory effect of NAc DBS on a global scale, differentially expressed gene (DEG) analysis was performed using Seurat^34^. Under adjusted P < 0.05□and□log2FC < |0.2| criteria, 104 up-regulated and 156 down-regulated DEGs were identified (Fig. 2A). Molecular function (MF) enrichment analysis for DEGs up-regulated under DBS showed enrichment of several Gene Ontology (GO) terms, such as ‘cell adhesion molecule binding’, ‘channel activity’, and ‘passive transmembrane transporter activity’ (Fig. 2B), and ‘catalytic activity, acting on RNA’, ‘methyltransferase activity’ and ‘transcription coregulator activity’ for down-regulated DEGs (Fig. 2C). Together with up- and down-regulated biological processes (BP) and cellular components (CC) enrichment terms (Supplementary Fig. S2A-D), our results implicated an upregulation in axon formation and connectivity, and a downregulation in protein synthesis and localization on a global scale. Examples of spatial expression pattern of up-(*Cmss1, Filip1l and Ttr*) and down-regulated (*Usmg5, Lcor and Gm43796*) DEGs showed an extensive and uniform distribution alteration (Fig. 2D). These results illustrated the global electrogenetic modulatory effect of NAc DBS.

**Fig. 2.**
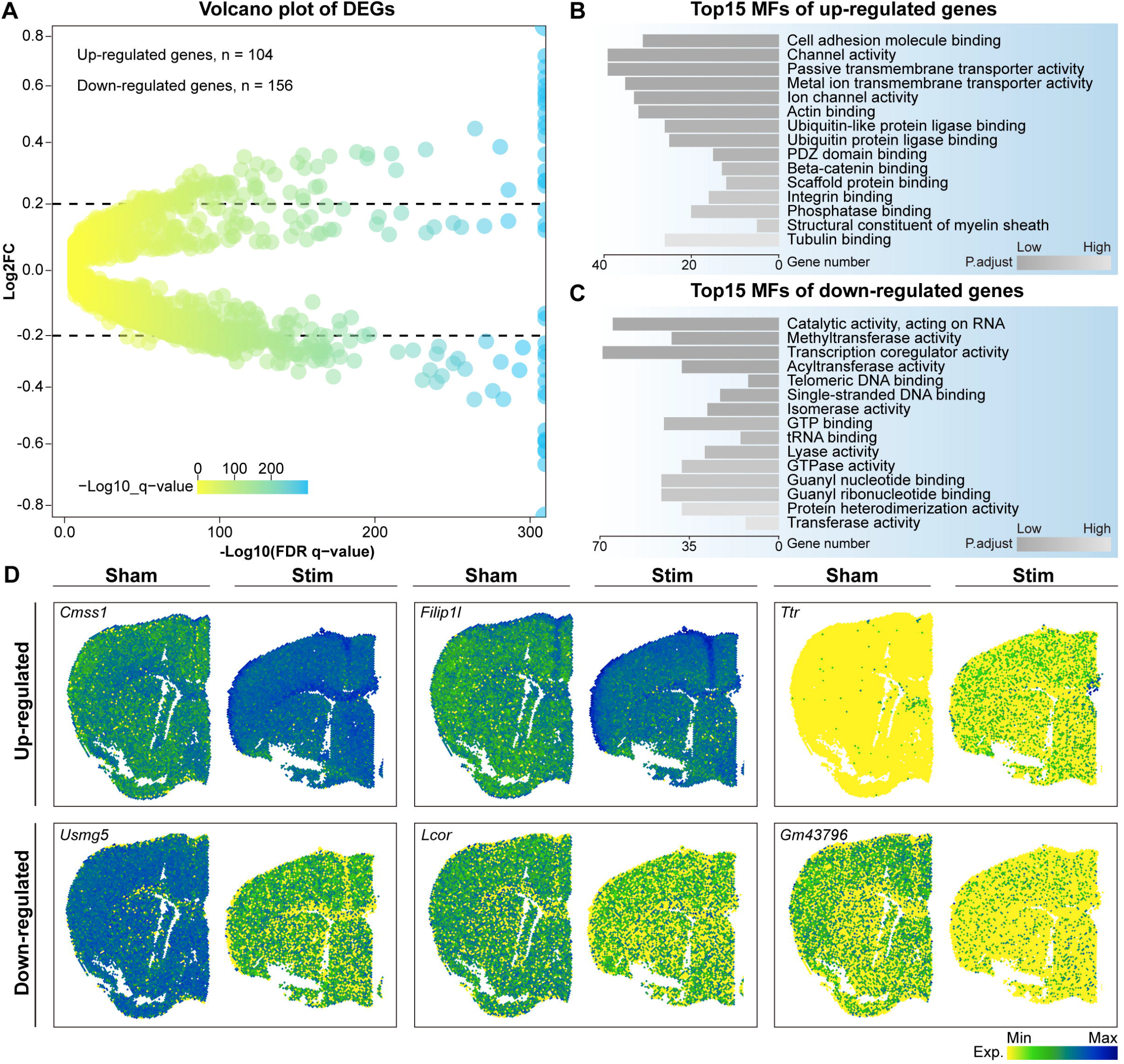
Effect of NAc DBS on global gene expression in the mouse brain. (**A**) Volcano plot showing differentially expressed genes (DEGs) between stimulation and sham-treated samples. Dots represent individual genes, with the color scale indicating the -log10 transformed adjusted p-value. Horizontal dotted lines represent a significance threshold, with a log2 fold-change threshold of |0.2|. (**B**) Top 15 Molecular Function (MF) enrichment results of up-regulated DEGs identified in (**A**). Enrichment analysis highlights pathways predominantly affected by up-regulated genes following deep brain stimulation (DBS) of the nucleus accumbens (NAc). (**C**) Top 15 Molecular Function (MF) enrichment results of down-regulated DEGs from (**A**), revealing pathways down-regulated by NAc DBS. (**D**) Spatial visualization of the expression of selected genes from (**A**) on Stereo-seq maps, illustrating the spatial distribution of significant gene expression changes in response to NAc DBS across the brain.

### DBS-induced modifications in the molecular architecture of the NAc neural circuits

After characterizing the cellular transcriptomic changes at the hemispheric level induced by NAc DBS, the next section of the survey explored alterations in neural circuits associated with the NAc. We executed DEG analysis and identified up- and down-regulated genes across different brain regions. NAc DBS induced region-specific changes in gene expression across multiple brain areas, as shown by the volcano plots (Fig. 3A).

**Fig. 3.**
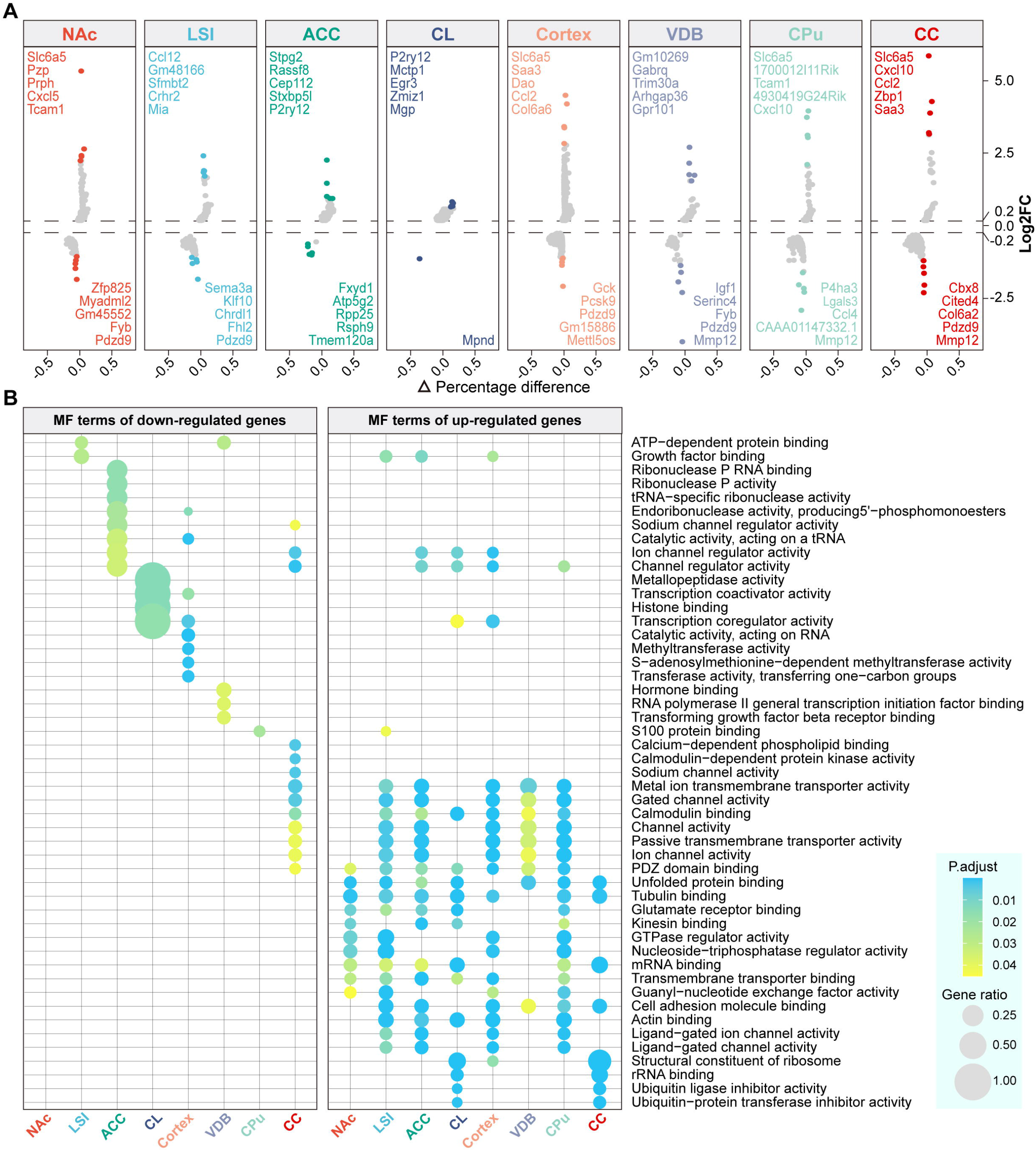
Electrical stimulation of NAc has differential impacts on gene expressions in various brain regions. (**A**) Volcano plots representing the differentially expressed genes (DEGs) across multiple brain regions. The top up-regulated and down-regulated DEGs are annotated at the top and bottom of each panel, respectively. The color gradient reflects the rank based on log2 fold-change, with a threshold of |0.2|. (**B**) Molecular Function (MF) enrichment analysis of up-regulated (right) and down-regulated (left) DEGs identified in (**A**). Dot size represents the gene ratio, which is the proportion of overlapped genes relative to the size of the Gene Ontology (GO) categories. The color scale indicates the adjusted p-value, reflecting the significance of the enrichment. CC (cingulate cortex), CPu (caudate putamen), LSI (lateral septal nucleus, intermediate part), NAc (nucleus accumbens), VDB (nucleus of the vertical limb of the diagonal band).

GO term analysis of the up-regulated DEGs revealed significant enrichment in pathways related to cell communication and signal transduction (Fig. 3B and Fig. S3A, right), including transmembrane transport, neuronal projection, and neurotransmitter release. We also observed that the up-regulated communication processes were predominantly localized to synaptic components (Fig. S3B, right). These findings confirmed that DBS not only altered synaptic function but also promoted structural reorganization of neuronal networks. Furthermore, spatial transcriptomics showed that this altered synaptic connectivity affected nearly every region examined, underscoring the broad impact of DBS in modulating connections within NAc-related circuits. Additionally, in CL and CC, we noted significant enrichment in pathways related to ribosome biogenesis and protein translation, suggesting a higher level of complexity and diversity in DBS-induced genetic effects at the circuit level.

In contrast to the relatively consistent enrichment pattern of up-regulated genes, the down-regulated GO terms were more structure-specific. In NAc and CC, the down-regulated terms were associated with pathways related to synaptic connectivity (Fig. S3A and Fig. S3B, left). In ACC, CL, and cortex, the down-regulated genes were enriched for pathways involved in DNA/RNA metabolism and processing (Fig. 3B and Fig. S3A, left). In CPu, GO terms related to stress responses, including regulation of inflammation, were significantly down-regulated (Fig. S3A, left). These results provide a more nuanced and detailed understanding of how DBS modulates both the target and related structures.

### Stereo-seq dissects NAc and identifies electrically altered genes at single-cell resolution

We then asked what electrogenetic effects DBS exerted on different cell types in the NAc. It has been assumed that, because of its large area size, spot of bin 100 contained multiple cell types. To enable single-cell gene expression profiling in the NAc, we performed DNA staining to highlight the nucleus, and then employed an image-based cell segmentation method, which can precisely integrate transcript and nucleic acid images into a gene-by-cell matrix (cell bin, Fig. 4A). Relative cell quantity in each cluster was measured (Fig. 4B). To eliminate the influence of brain section sizes, we calculated the absolute number of cells from each cluster corrected by total NAc bin 100 counts (Fig. 4C). In the stimulated samples, neurons, including D1-receptor-, D2-receptor-, AMPAR- and NMDAR-expressing neurons, showed a marked decrease in density compared to sham-treated samples, while other cell types, such as interneurons, astrocytes and microglia, exhibited smaller changes. This suggests that DBS selectively reduces the density of GABAergic neurons in the NAc.

**Figure 4.**
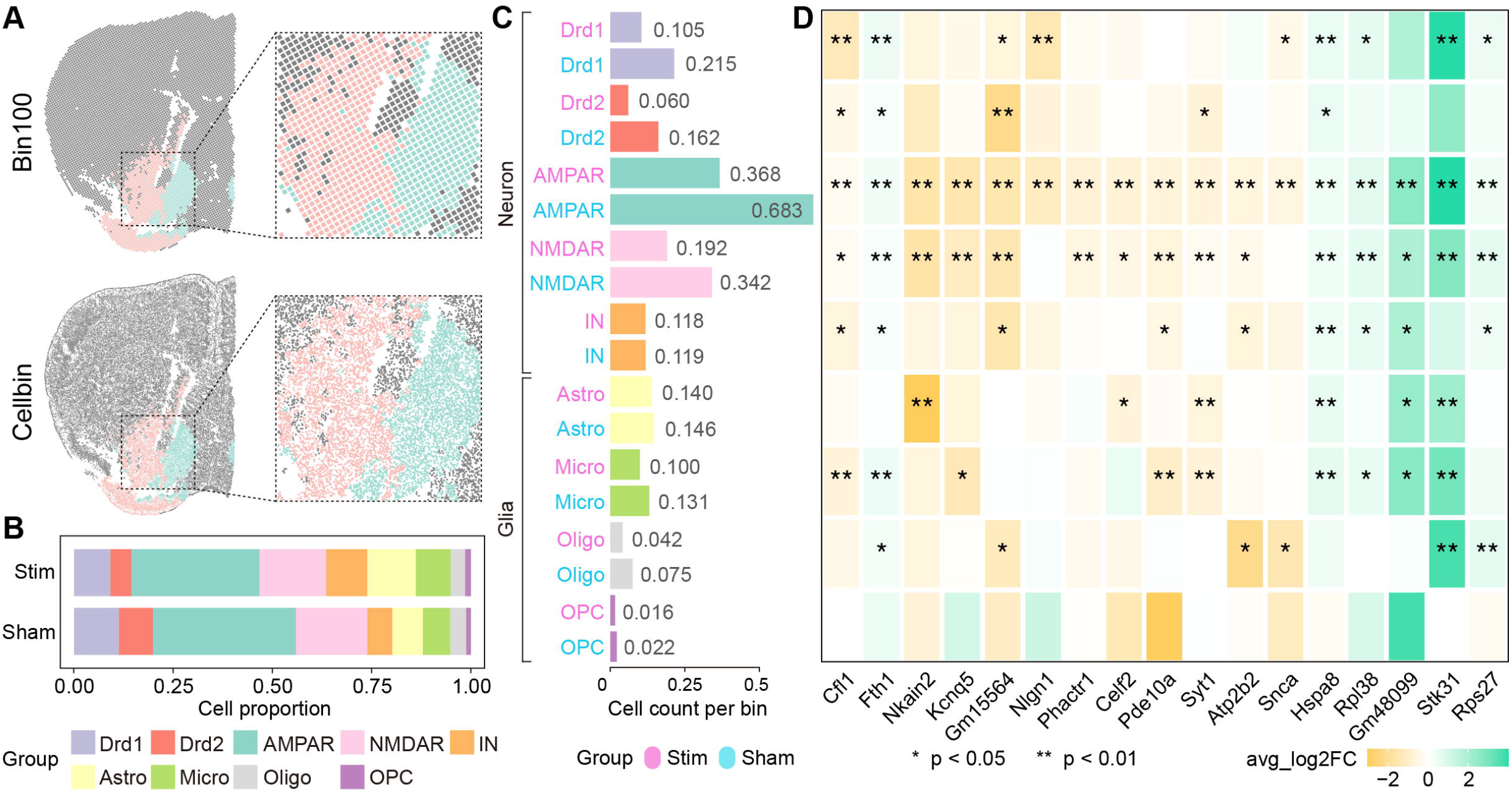
Stereo-seq dissects differential cell type-specific responses to electrical stimulation in the NAc. **(A)** Schematic representation of spatial visualizations using bin100 (upper) and cellbin (lower) methods. The insets show magnified views of the core (red) and shell (green) of the nucleus accumbens (NAc). **(B)** Proportional cell type composition in the stimulated (Stim) and sham-treated (Sham) groups, including Drd1 (dopamine receptor D1-expressing neurons, purple), Drd2 (dopamine receptor D2-expressing neurons, red), AMPAR (AMPA receptor-expressing neurons, teal), NMDAR (NMDA receptor-expressing neurons, pink), interneurons (orange), astrocytes (yellow), microglia (green), oligodendrocytes (light gray), and oligodendrocyte precursor cells (OPCs, dark gray). Bar plots represent the relative abundance of each cell type in the two conditions. **(C)** Cell count per bin100 for each cell type in the stimulated and sham groups. The counts reflect the number of cells per bin across the different cell types, with Drd1-, D2d2-, AMPAR-, and NMDAR-expressing neurons showing significant decrease after electrical stimulation. **(D)** Heatmap of average log2 fold change (log2FC) in gene expression across key genes in response to NAc DBS. The color gradient represents the average log2FC, with green indicating up-regulation and yellow/orange indicating down-regulation. Genes such as Nkain2, Snca, Pde10a, and Stk31 were significantly down-regulated in stimulated samples, whereas genes such as Stk31 and Hspa8 were up-regulated in response to stimulation. Interestingly, most of these differentially expressed genes (DEGs) were found within the AMPAR (17 out of 17) and NMDAR (15 out of 17) clusters, suggesting that NAc DBS primarily influences glutamate receptor-expressing neurons through its electrogenetic effects. Asterisks denote the level of statistical significance (*: p-value ≤ 0.05; ** : p-value ≤ 0.01).

In-depth analysis of the spatial transcriptomics data revealed that DBS in the NAc region elicited differential gene expression patterns across distinct neuronal and glial cell types (Fig. 4D). The identities of each cell bin was determined based on established cell marker genes, such as dopamine D1 receptor marker Drd1, dopamine D2 recepor marker Drd2, AMPA receptor marker Gria2, NMDA receptor marker Grin1, interneuron marker Resp18, astrocyte marker Gja1, microglia marker C1qa, oligodendrocyte marker Mog and oligodendrocyte precursor cells marker Pdgfra. Under adjusted P < 0.05 threshold, we identified a total of 17 DEGs across cell types (mean expression levels and fold changes in expression of these genes were displayed in Fig. 4D and Supplementary Fig. S4, respectively). Key genes involved in synaptic transmission and plasticity, including neuroligin-1 encoding gene Nlgn1 (crucial for synapse formation and function in neurons), synaptotagmin-1 encoding gene Syt1 (essential for calcium-ion-triggered neurotransmitter release), alpha-synuclein encoding gene Snca (regulates synaptic vesicle trafficking and subsequent neurotransmitter release), Na_+_/K_+_-ATPase-interacting protein encoding gene Nkain2 (potentially involved in neuronal ion transport), voltage-gated potassium channel encoding gene kcnq5 (contributing to neuronal excitability), and phosphodiesterase 10A encoding gene PDE10a (an enzyme involved in neuronal signal transduction by hydrolyzing cyclic nucleotides), were down-regulated after NAc DBS compared to sham stimulation. Other genes modulated by NAc DBS include down-regulated Cofilin-1 and Phactr1 (cytoskeletal), Celf2 (RNA binding), Fth1 (iron storage), Atp2b2 (calcium pump), and up-regulated Hspa8 (heat shock protein stress responses), Stk31 (serine/threonine kinase), and Rpl38 and Rps27 (ribosome). Remarkably, the majority of these DEGs were found in AMPAR (17 out of 17) and NMDAR (15 out of 17) clusters, which indicated that NAc DBS exert an electrogenetic effect predominantly on glutamate-receptors-expressing neurons. Additionally, glial cells, including astrocytes and microglia, also exhibited changes in gene expression. Genes involved in cellular stress responses, such as Hspa8 and Stk31, showed differential expression patterns, suggesting that DBS may exert neuroprotective effects by creating a more resilient microenvironment.

Overall, these findings demonstrate that DBS of the NAc leads to cell type-specific alterations in gene expression, and provide key insights into the molecular mechanisms underlying the therapeutic effects of DBS in modulating brain activity. Based on these findings, a putative cellular mechanism of NAc DBS for neuropsychiatric disorders is proposed and summarized in Fig. 5.

**Fig. 5.**
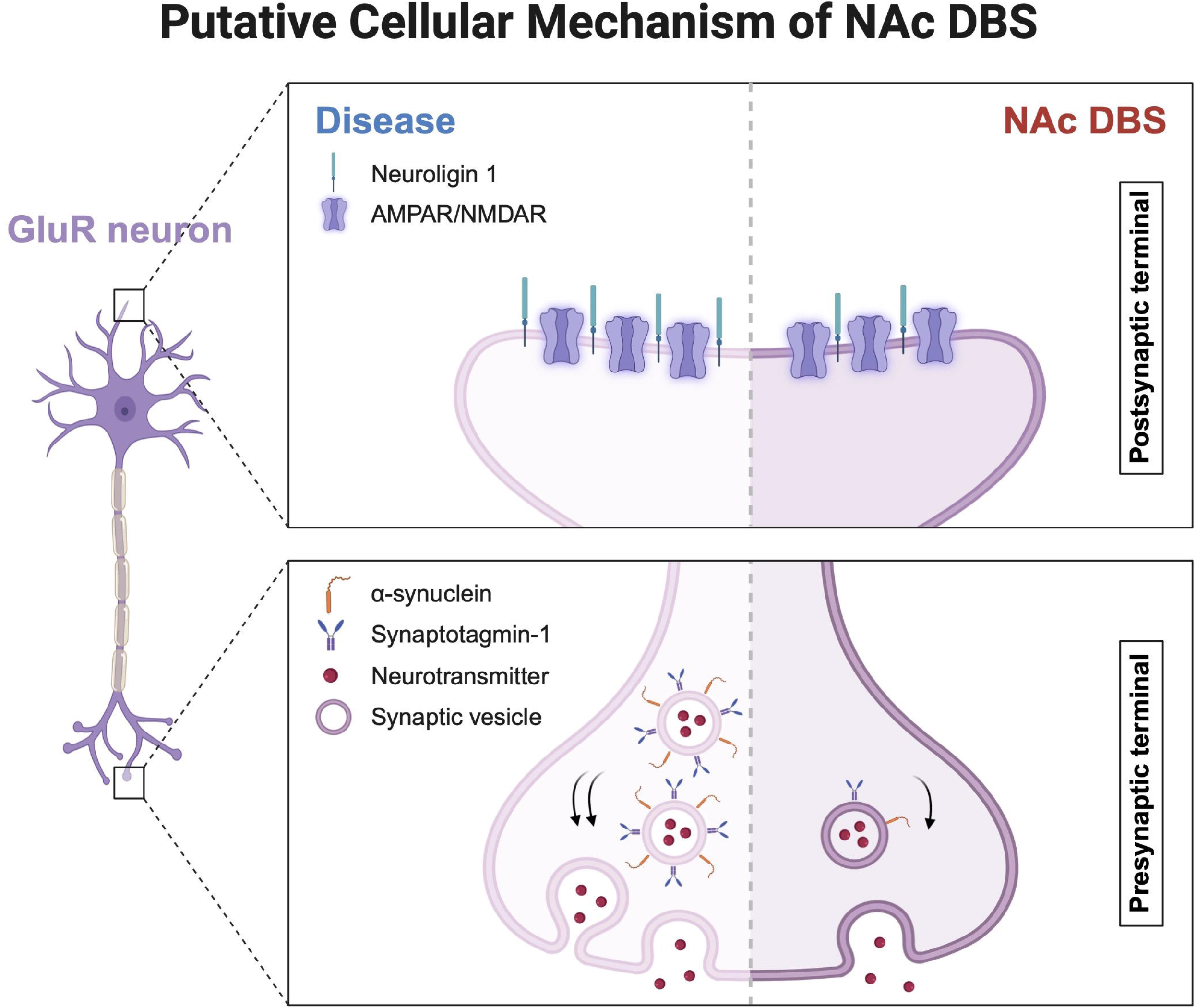
Putative therapeutic mechanisms of NAc DBS in glutamate receptor expressing neurons. Schematic diagram of a glutamate receptor expressing neuron. Presynaptic and postsynaptic areas are magnified in right panels (top and bottom, respectively). In the postsynaptic terminal, compared to disease states (left), NAc DBS reduces the expression of neuroligin 1 to modulate neurotransmission and neuroplasticity (right). In the presynaptic terminal, NAc DBS reduces the expression of proteins involved in synaptic vesicle trafficking, including α-synuclein and synaptotagmin-1, which regulate synaptic vesicle exocytosis and neurotransmitter release. This leads to decreased neurotransmitter release and reduced communication between neurons. Arrows indicate the direction of synaptic vesicle movement and neurotransmitter release. This figure was created using BioRender platform (www.biorender.com).

## DISCUSSION

Deep brain stimulation (DBS) is a promising neuromodulation intervention for treatment-resistant neuropsychiatric disorders; however, its mechanisms of action are not yet fully understood. In our study, we present the first use of high-resolution spatial transcriptomics to explore the molecular mechanisms underlying nucleus accumbens (NAc) DBS. Our findings reveal that NAc DBS induces widespread alterations in gene expression on a global scale. Additionally, we observed changes in gene expression pointing toward remodeled synaptic connectivity in the NAc circuit, which is implicated in various neuropsychiatric conditions. Notably, the DBS-induced synaptic adaptations exhibit a bidirectional pattern, engaging both upstream and downstream structures of the NAc. For instance, the NAc receives direct inputs from ACC projections, and this ACC-NAc circuit integrates both reward and effort information^37^. Previous studies reported that GABA deficit in ACC correlates specifically with major depressive disorder (MDD) in adolescents^38, 39^. We found that in ACC, Cep112 is one of the top-5 most significant upregulated DEGs by NAc DBS, and the protein encoded by this gene is thought to interact with GABA receptors in neurons^40^. In addition to GABA, another critical finding was the gene enrichment in GO temrs of glutamate receptor binding in regions connected to the NAc, including the ACC. There is evidence that glutamate concentration in the ACC is downregulated under neuropsychiatric state^41^. Our findings indicate that the upregulated glutamate receptor binding in the ACC may reverse the dysregulation of glutamate. LSI is another upstream brain structure of NAc known to play a critical role in the reward circuit^42^. Sema3a gene was found down-regulated in LSI after NAc DBS, whose pathway is involved in the localization of glutamate receptor^43^. This downregulation of glutamate transmission may help restore balance in the reward circuit^44^. These observations align with previous studies indicating that electrical stimulation can trigger local action potentials propagating both orthodromically and antidromically, eliciting synaptic events and exerting effects to induce molecular remodeling at connected areas^45-48^.

Such synaptic network remodeling involves multiple stages of plasticity. By examining both up- and down-regulated GO terms, we identify molecules associated with axonogenesis, transmembrane transport, adaptor and scaffolding proteins, cytoskeletal elements, and signaling molecules in the NAc circuit. These processes are essential for neuronal communication and downstream signal transduction, forming a linked chain of events that modify synaptic plasticity at both pre- and postsynaptic levels. Disruptions in the regulation of synaptic transmission and neuronal activity are commonly associated with neuropsychiatric disorders^49-53^. Accumulating evidence suggests that DBS promotes synaptic plasticity and network reorganization^54-56^. NAc DBS may restore disrupted synaptic signaling pathways through fine-tuned regulation, thereby exerting its therapeutic effects in neuropsychiatric conditions.

Supporting this notion, the 17 differentially expressed genes (DEGs) identified in the NAc are primarily involved in synaptic transmission. Their specific functions range from synapse formation (Nlgn1), neurotransmitter release (Snca and Syt1), ion channel regulation (Nkain2 and Kcnq5), to signal transduction (PDE10a). These genes are associated with various neuropsychiatric disorders. For example, inhibition of PDE10a, which is selectively expressed in the striatum including the NAc, has been proposed as a treatment for opioid addiction^57^. It has also been shown that PDE10A plays a crucial role in integrating the functions of D1 and D2 MSNs^58^. Nlgn1 deletion or duplication is linked to obsessive-compulsive disorder^59^, while Snca overexpression, which impairs synaptic function, has been associated with depressive-like behaviors and positively correlates with depression severity^60, 61^. Syt1 is crucial for the long-term regulation of alcohol dependence^62^. Nkain2 has been implicated in bipolar disorder, alcohol addiction, and psychiatric traits^63-65^, and KCNQ5 potassium channel modulators have emerged as promising supportive therapies for major depressive disorder^66^. Our findings indicate by downregulating these genes, DBS attenuates synaptic activity and plasticity and modulates the hyperactive neural circuits implicated in neuropsychiatric disorders.

Some of the identified genes may interact with other risk-related components. For example, Phact1, an actin-binding protein, regulates Slack channels, which in turn modulate anxiety-like behaviors^67, 68^. Taken together, these genes provide insights into how NAc DBS may bridge the gap between its therapeutic effects and the underlying molecular mechanisms. Our findings suggest that DBS may reverse the synaptic transmission dysfunction associated with neuropsychiatric disorders by targeting these genes involved in synaptic process.

Further analysis revealed that AMPAR- and NMDAR-expressing neurons exhibit the greatest convergence of identified DEGs, suggesting that these neurons may have the strongest influence on the effects of NAc DBS. AMPAR- and NMDAR-dependent synaptic plasticity within the NAc is crucial in reward-seeking behaviors^69, 70^. These neurons are likely affected by glutamatergic inputs from upstream structures such as the amygdala, prefrontal cortex (PFC), and ventral tegmental area (VTA)^71-73^. Prior study has shown that PFC and VTA glutamatergic projections to the NAc are critical for regulating reward and drug-seeking behaviors^74, 75^. DBS has been found to significantly reduce glutamate levels in both the NAc and VTA, indicating that NAc-DBS inhibits morphine-related hyperactivation of neurotransmission within the reward circuit^76^. Consequently, a cascade of molecular events is likely to occur in glutamate receptor-expressing neurons^77^. A possible explanation for this highly specific electrogenetic modulation is the resetting of the voltage-gated component of glutamate receptors after DBS^78^. NMDAR-dependent plasticity is thought to induce long-term potentiation as well as long-term depression at different synapses^79^, which may help normalize the neuronal transmission manifested in neuropsychiatric conditions. AMPAR and NMDAR blockers also show antidepressant and anti-addiction effects^80^, in alignment with our findings. This work demonstrates the importance of glutamate receptor neurons in the action mechanism of NAc DBS, and provide specific molecular targets for further investigation.

We also identify the possible involvement of non-neuronal cells in the modulatory effects of DBS. In the caudate putamen (dorsal striatum), we observe a downregulation in processes related to the negative regulation of the immune system, such as T-cell activation and leukocyte cell-cell adhesion. Dysregulation of the immune system has been implicated in various neuropsychiatric disorders, including depression and OCD^81, 82^. Our findings, aligned with previous studies, suggest that NAc DBS may modulate the immune system, potentially influencing reward-related processes in the caudate putamen^83-85^.

One limitation of our study is the relatively small sample size. Future research involving larger cohorts and additional brain regions will be crucial to validate and extend our findings. Combining spatial transcriptomics with electrophysiological, imaging, and behavioral assessments would provide more robust evidence regarding the mechanisms underlying DBS. Single-cell sequencing could further highlight the significance of the identified differentially expressed genes. Moreover, employing targeted disease models could reveal additional gene changes specific to individual conditions, while testing these findings in human samples will be essential to confirm their translational potential.

This study advances previous research by offering a high-resolution view of the molecular changes induced by DBS. Prior investigations of DBS’s genetic modulation were limited by lower spatial resolution and focused on narrow gene sets^25, 86, 87^. Our findings are consistent with a previous RNA-sequencing study, which also reported that DBS-induced gene changes promote neuroplasticity^25^. By employing Stereo-seq technology, we mapped the spatial distribution of gene expression changes at near single-cell resolution, providing a more comprehensive view of the global and region-specific molecular effects of DBS. Our study builds on prior work by showing that DBS modulates a wide range of genes and cellular processes, primarily related to synaptic transmission, both at the stimulated site and in connected brain areas. These results have important implications for the clinical application of DBS. By identifying specific genes and pathways affected by DBS, we highlight potential pharmacological targets that could enhance the therapeutic efficacy of DBS. For example, drugs that modulate cell adhesion molecules or glutamate receptor activity might be used in conjunction with DBS to improve treatment outcomes for patients with treatment-resistant neuropsychiatric disorders. Additionally, our approach opens new avenues for optimizing DBS parameters, such as stimulation intensity and frequency, by providing a molecular blueprint based on detailed gene expression mapping.

In conclusion, our research sheds light on the dual role of NAc DBS in influencing synaptic and neuroplastic processes while mitigating stress and inflammatory responses in targeted brain circuits. The upregulation of genes involved in synaptic functions and the downregulation of those related to stress responses underscore the therapeutic potential of DBS in neuropsychiatric disorders. These findings provide a molecular basis for how DBS modulates neural circuits, offering novel insight into its mechanisms and guiding future strategies to refine and optimize DBS treatments. Further studies should focus on validating these molecular insights across broader populations and refining clinical applications to enhance therapeutic outcomes for patients with resistant neuropsychiatric conditions.

## DATA AVAILABILITY

The raw data generated in this study have been deposited to CNGB Nucleotide Sequence Archive with accession number CNP0006277 (https://db.cngb.org/search/project/CNP0006277). The code used to generate the results of this study is available on the GitHub repository at https://github.com/Hemmings-Wu-Lab/NAc-DBS-Transcriptomics.

## ACKNOWLEDGEMENTS

We thank Chang Wang for her contribution in experimental procedure in this study.

## AUTHOR CONTRIBUTIONS

CC, HW, and JZ conceived and designed the project. CC, LG and HW contributed to the analysis and interpretation of data and wrote the manuscript. CC and ZZ contributed to the data acquisition, statistical analyses and prepared the tables and figures. LG, WC, CY and KX verified the results. LG, HW and JZ edited the manuscript and provided supervision. All authors read and approved the final manuscript.

## ETHICS APPROVAL

All animal procedures were approved by the second affiliated hospital of Zhejiang university – Animal Ethics Committee (Approval No. AIRB-2021-087) and were performed according to the Guiding Principles for the Care and Use of Laboratory Animals Approved by Animal Regulations of National Science and Technology Committee of China.

