## Supplementary figures and images for "Change in brain molecular landscapes following electrical stimulation of the nucleus accumbens"

### Figure S1

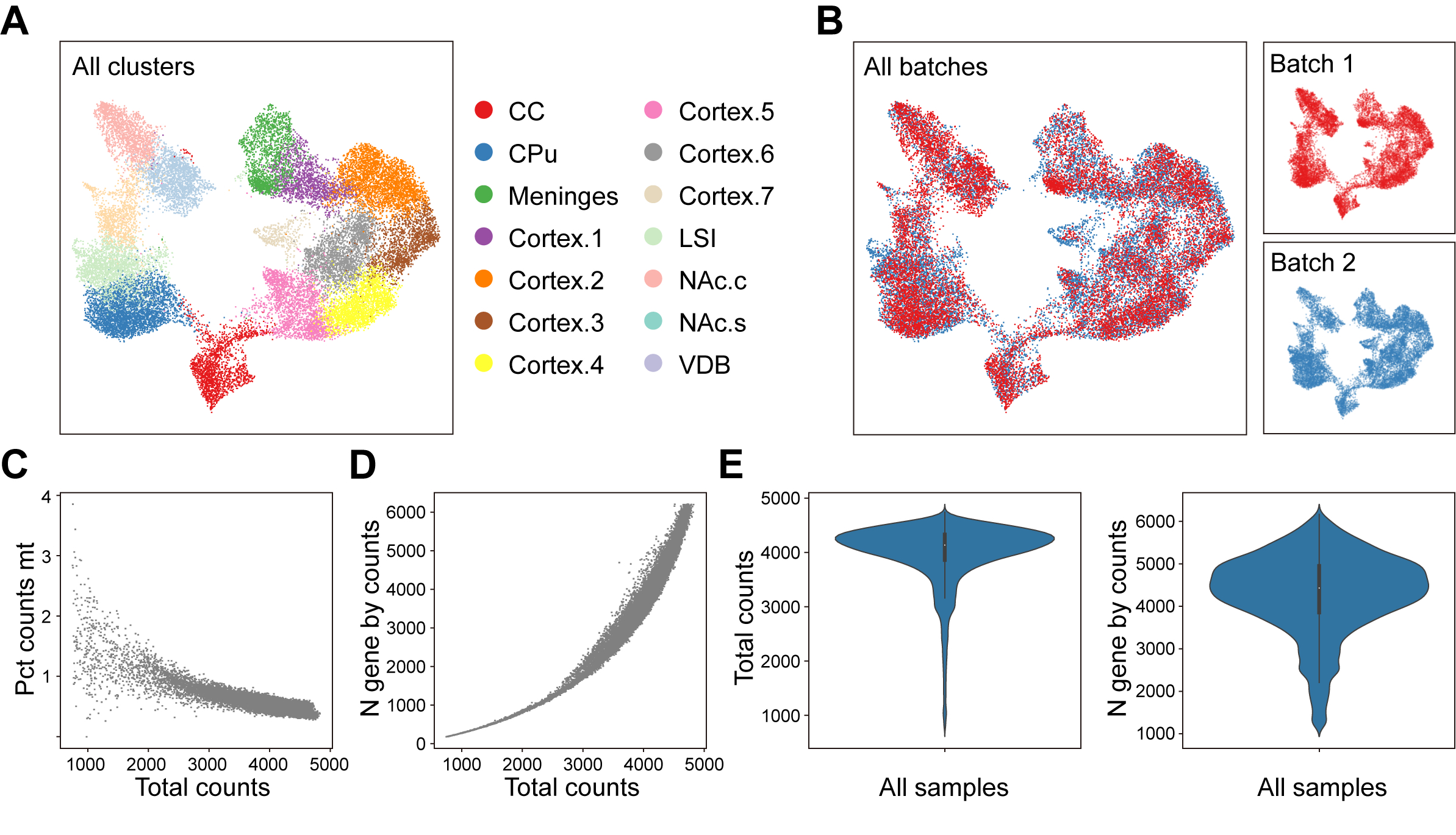

### Figure S2

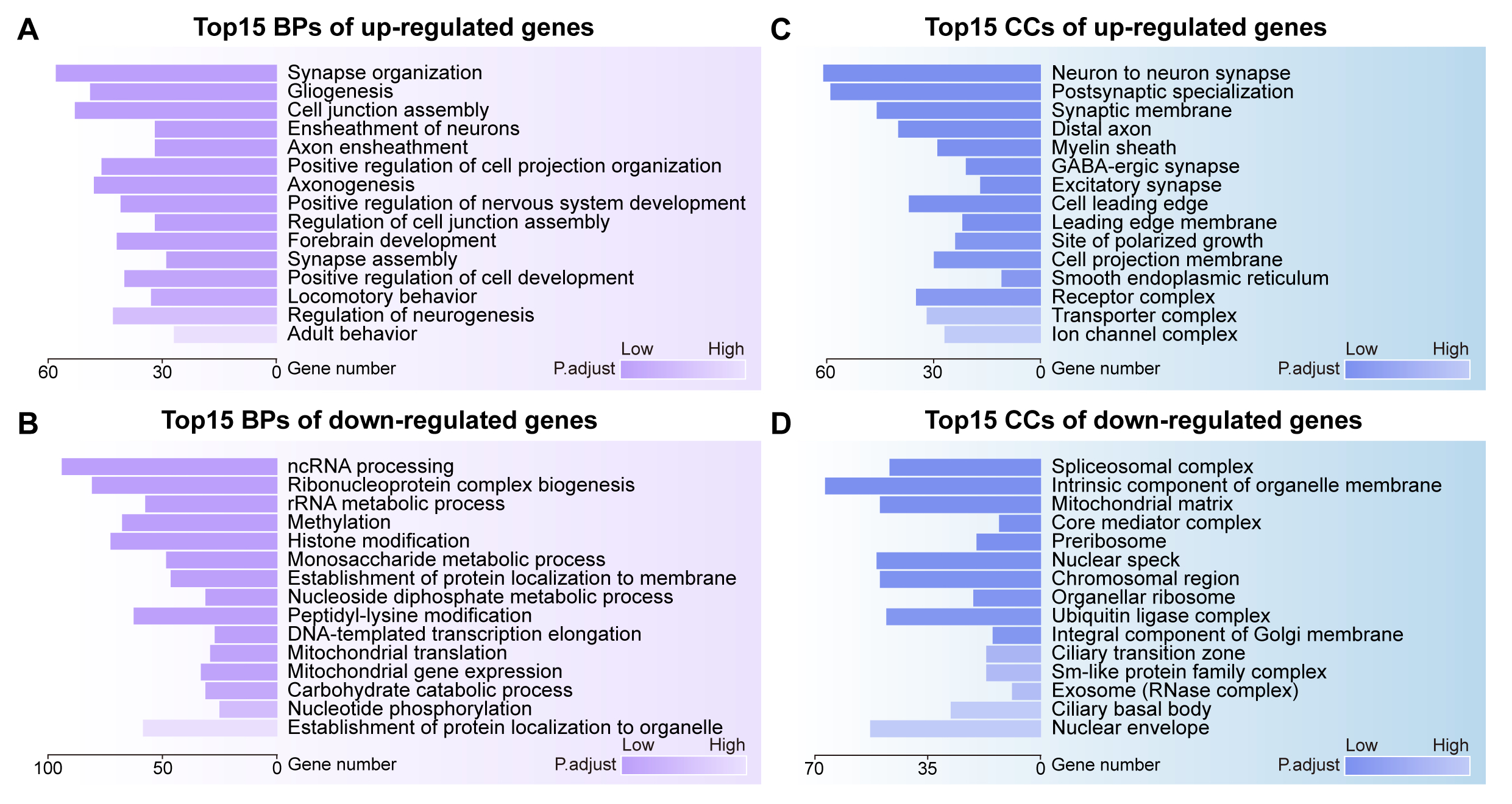

### Figure S3

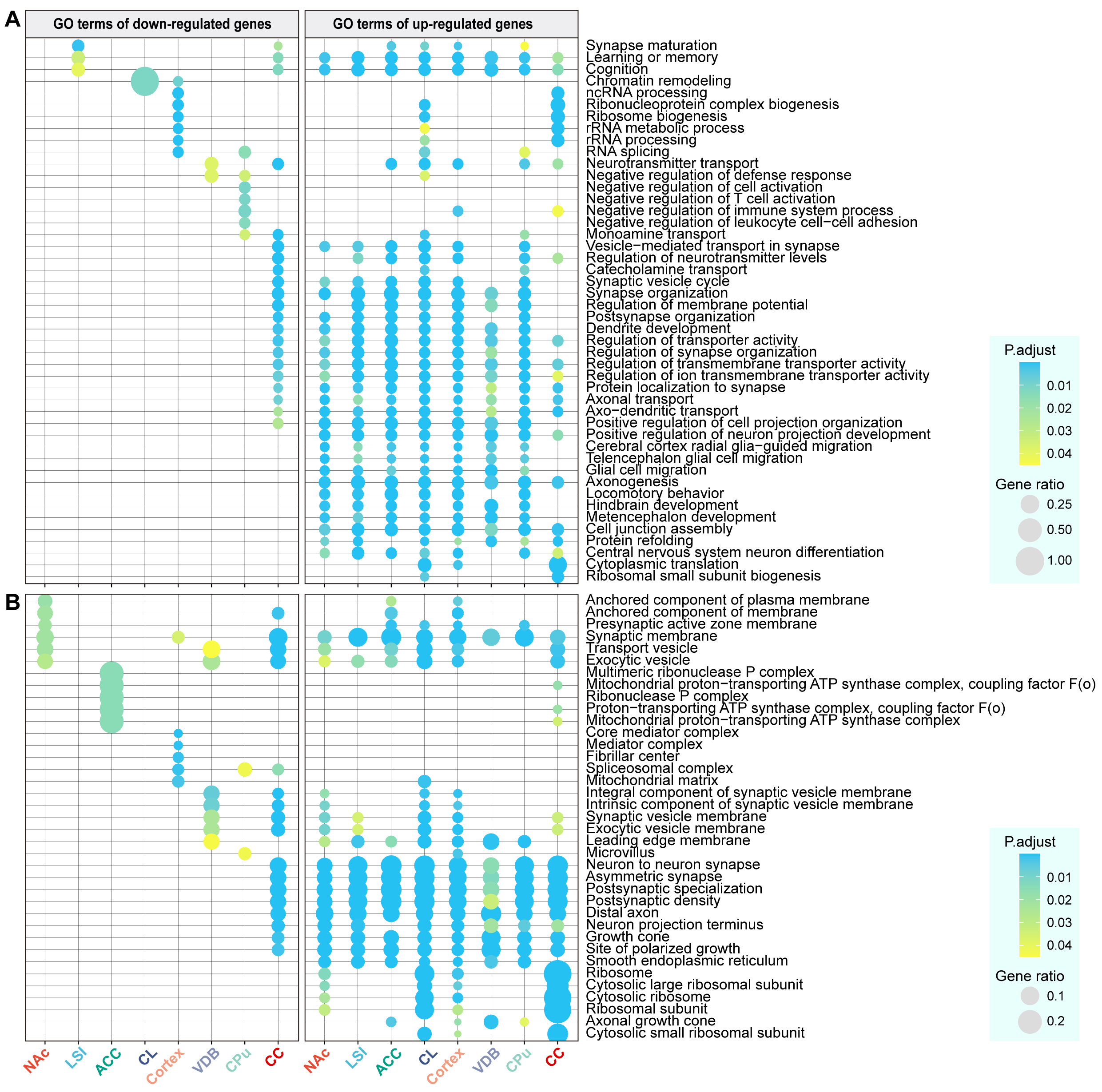

### Figure S4

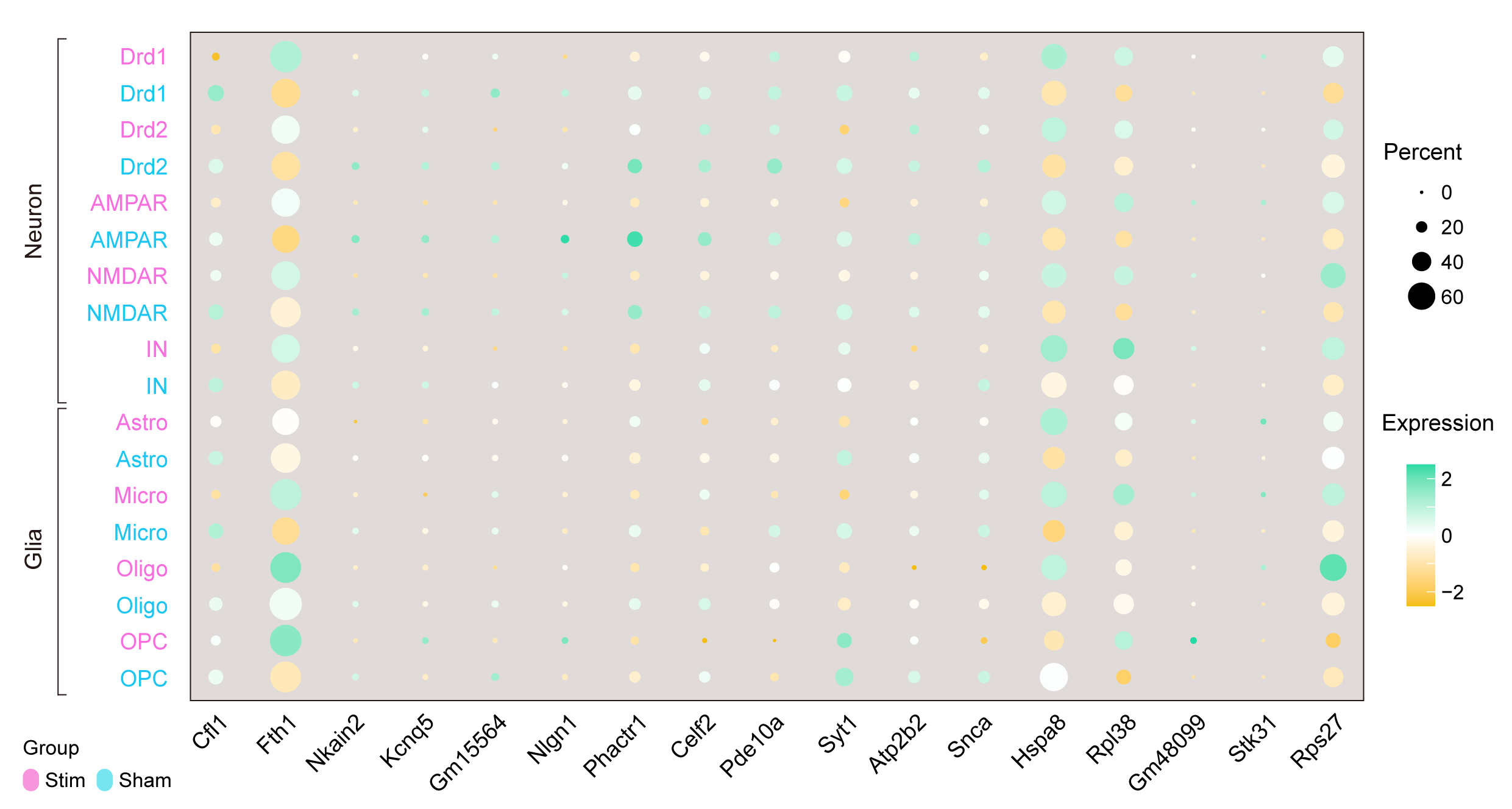
