## Supplementary information for "Change in brain molecular landscapes following electrical stimulation of the nucleus accumbens"

**Supplementary figures legends**

**Figure S****1.** **Performance of stereo-seq****, related to Fig. 1.** (**A**) UMAP plot displaying identified clusters in Fig. 1B. (**B**) UMAP plots of the different batches of samples. (**C**) Scatter plot showing the correlation between total UMI counts and percentage of mitochondrial gene counts. (**D**) Scatter plot showing the correlation between total UMI counts and gene numbers (right). (**E**) Violin plots of UMI count (left) and gene count (right) for overall samples. CC: Cingulate cortex; CPu: Caudate putamen; LSI: intermediate part of the lateral septal nucleus; NAc.c: Nucleus accumbens, core; NAc.s: Nuclues accumbens, shell; VDB: nucleus of the vertical limb of the diagonal band.

**Figure S2.** **Functional enrichment analysis results of global DEGs, related to Fig 2.**

**2.** (**A**) The top 15 BP enrichment results of up-regulated DEGs from Fig. 2A. (**B**) The top 15 BP enrichment results of down-regulated DEGs from Fig. 2A. (**C**) The top 15 CC enrichment results of up-regulated DEGs from Fig. 2A. (**D**) The top 15 CC enrichment results of down-regulated DEGs from Fig. 2A.

**Figure S3. Functional enrichment analysis results of DEGs** **in different brain regions, related to** **Fig. 3.** (**A** and **B**) BP (A) and CC (B) enrichment results of up- (right) and down-regulated (left) DEGs in Fig. 3A. The size of the dot represents gene ratio between overlapped genes and GO categories size, and the color represents the adjusted p-value. ACC: Anterior cingulate cortex; CC: Cingulate cortex; CL: Claustrum; CPu: Caudate putamen; LSI: Intermediate part of the lateral septal nucleus; NAc.c: Nucleus accumbens, core; NAc.s: Nuclues accumbens, shell; VDB: nucleus of the vertical limb of the diagonal band.

**Figure S4. Dotplot of DEGs across cell types in the nucleus accumbens, related to Fig. 4.** The color of dots indicates average expression, and the size of dots represents average percent of cells expressing selected gene.
